# Older is Order: Entropy reduction in cortical spontaneous activity marks healthy aging

**DOI:** 10.1101/2024.11.13.623510

**Authors:** Da Chang, Xiu Wang, Yaojing Chen, Zhuo Han, Bing Liu, Zhanjun Zhang, Xi-Nian Zuo

## Abstract

Entropy trajectories remain unclear for the aging process of human brain system due to the lacking of longitudinal neuroimaging resource. We used open data from an accelerated longitudinal cohort (PREVENT-AD) that included 24 healthy aging participants followed by 4 years with 5 visits per participant to establish cortical entropy aging curves and distinguish with the effects of age and cohort. This reveals that global cortical entropy decreased with aging, while a significant cohort effect was detectable that people who were born earlier showed higher cortical entropy. Such entropy reductions were also evident for large-scale cortical networks, although with different rates of reduction for different networks. Specifically, the primary and intermediate networks reduce their entropy faster than the higher-order association networks. We conclude two specific characteristics of the entropy of the human cortex with aging: the shift of the complexity hierarchy and the diversity of complexity strengthen.

## 1 Introduction

Understanding the brain mechanism of aging progression has been a major concern in successful aging. As a living dynamic system, the human brain processes information with high complexity. However, we still know little about this complexity and its aging patterns. Entropy is a concept that ranges from thermodynamics to complexity of a physical dynamic system. Higher entropy indicates a more complex system with more irregularity [1] or uncertainty [2] of its dynamics. Unlike increasing monotonically over time in a closed system (e.g., the universe), entropy remains relatively low in a biological system by continually exchanging energy with environment to maintain its own orderliness. In other words, a living organism feeds upon extracting ‘order’ from the environment, a stream of ‘negative entropy (NE)’, to compensate the increases of intrinsic entropy for maintaining its organization [3]. The human brain, as a complex living system, thus should operates on the same pattern. Its function becomes more ordered and predictable with the accumulation of more experience and knowledge. The entropy of the human resting state brain has been found to be negatively correlated with education [4, 5] and task performance [4] and, more directly, reduced due to periodic participation in tasks [6].

Aging is normally associated with greater predictability or less complexity of neurophysiological processes [7–9]. In line with the NE pattern, we propose a hypothesis that the aging progression of human brain function is marked by entropy reduction during normal aging. Interestingly, modern neuroimaging studies did not converge into this prediction while demonstrating that entropy decreases [7, 10–15], increases [5, 11], or does not change [6] with age. These inconsistent observations may reflect the cross-sectional nature of the experimental design in previous studies. The cross-sectional method has been well developed for investigations of individual differences. Cross-sectional changes related to age have been validated as related to interindividual differences in early life factors but not to longitudinal brain change [16], that is, aging progression. A direct examination of longitudinal entropy changes in the proposed hypothesis of aging progression is missing, while the other assessments of aging progression on human brain function are also very rare due to limited longitudinal data and imaging methodology (see an exception in [17]).

Functional magnetic resonance imaging (fMRI) can detect signals dependent on the blood oxygen level (BOLD) in vivo to measure spontaneous brain activity during the resting state [18–20], and thus makes large-scale longitudinal data collection convenient. The sample entropy (SampEn) is a nonparametric entropy metric to measure the complexity of fMRI and other physiological signals [21, 22]. The lower SampEn indicates more predictable or less complex brain dynamics, while the higher SampEn indicates less predictable or more complex brain dynamics. This entropy metric has been validated as a reliable measure of temporal brain dynamics [4, 6] and exhibited promising prediction validity in discovering entropy of brain functions [10, 23–28]. Here, we used longitudinal fMRI data from the PREVENT-AD (PResymptomatic EValuation of Experimental or Novel Treatments for AD) database [29, 30] to chart the entropy trajectories of the human brain cortex during the progression of aging using SampEn. We expect our hypothesis that entropy reduces during aging to be confirmed for both whole cortex and large-scale networks, while such NE pattern is differentiable across different networks according to their hierarchical orders of functional organization.

## 2 Results

### 2.1 The PREVENT-AD sample

We used an accelerated longitudinal sample from the PREVENT-AD cohort. It consists of 24 Canadian participants who received 5 annual visits, including 1 baseline visit and 4 follow-up visits (the 12, 24, 36, 48 months after the baseline visit). These individuals without cognitive impairment were recruited from families with a history of AD in their parents or multiple siblings (16 females, age range at the baseline visit: 58-77 years old, age cover with the full sample: 58-82 years old). The frequency of APOE-*ϵ*4 in this sample is 12.5%, which is comparable to the frequency of typical Caucasian populations (15%)[31]. All participants completed structural magnetic resonance imaging (MRI) and fMRI, as well as cognitive tests, which are documented in detail by [29].

### 2.2 Global entropy reduction

Following our hypothesis, the global cortical SampEn decreased during the aging progression. This aging effect was contaminated with a cohort effect while gene, sex and education had no effects on the global entropy. Specifically, the generalized additive mixed model (GAMM) revealed an adjusted *R*^2^ of 16.9%, explaining the data variability by both the aging and the cohort effect. In Figure 1, each segment of the heavy blue line depicts the aging curve of individuals from a specific age cohort, indicating consistent linear reduction patterns in different cohorts (*p* = 0.0073). The cross-sectional age trajectory showing slight U-shaped changes in SampEn concealed this aging process at the population level (the heavy blue dashed line) with the cohort effect (*p* = 0.0263, the red dashed line), which showed that the earlier-born cohorts exhibited significantly higher cortical SampEn relative to the later-born cohorts.

**Fig. 1.**
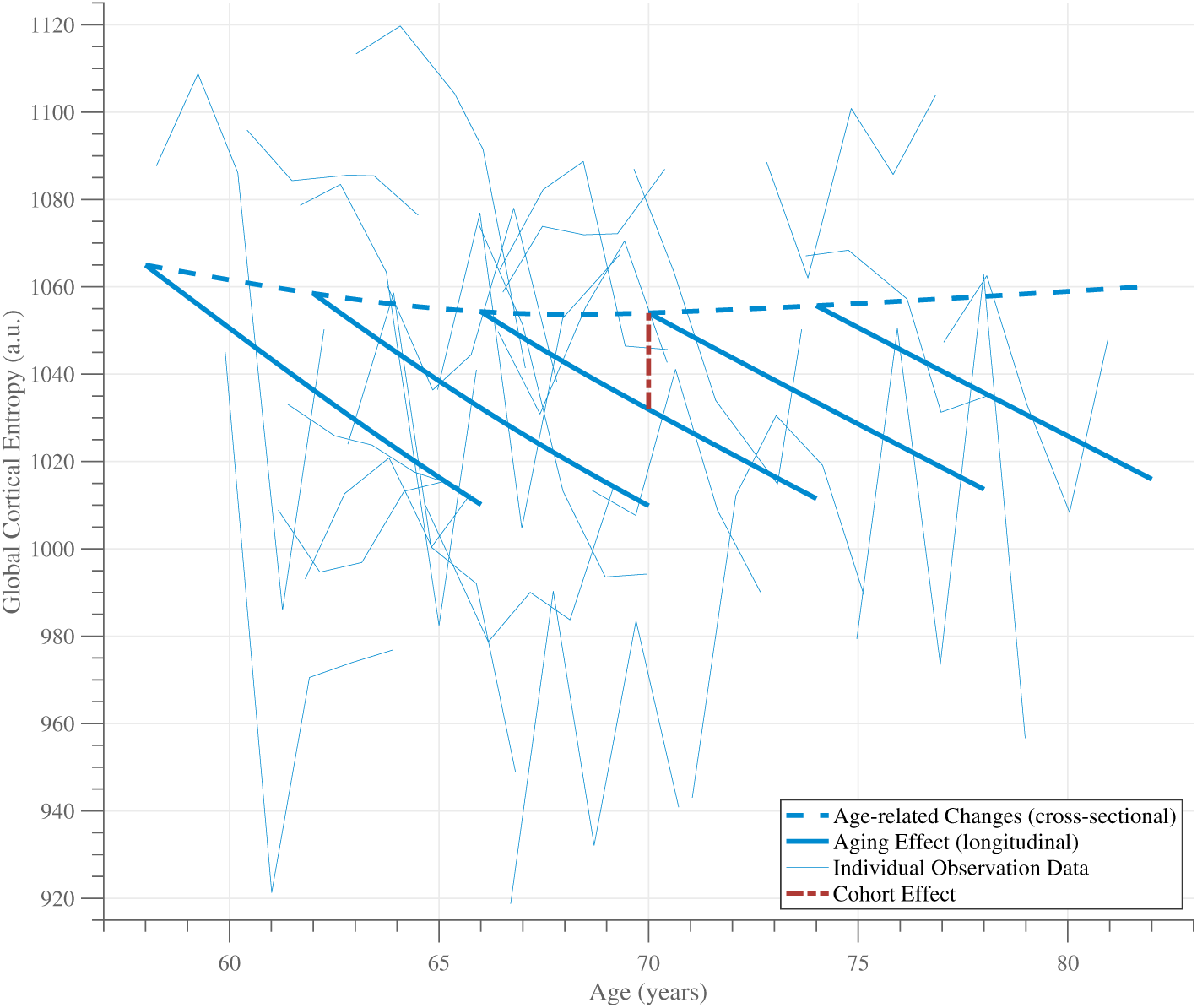
Aging pattern of global entropy in the human cerebral cortex. Five repeated measures of the global SampEn for each participant are connected into individual background blue thin lines. Model fitted by GAMM generated the heavy blue dash line and blue line segments. These line segments extending from the blue dash line indicate the within-subject changes from five different age cohorts (58, 62, 66, 70, and 74 years of age at baseline), the aging curves. The dashed blue curve represents age-related changes of global entropy while the dashed red segment indicates the cohort effect.

### 2.3 Network entropy reduction

The fifteen functional networks of the human cerebral cortex are used to examine the entropy aging patterns at network level: Visual-Central (VIS-C), Visual-Peripheral (VIS-P), Somatomotor-A (SMOT-A), Somatomotor-B (SMOT-B), Auditory (AUD), Premotor-Posterior Parietal Rostral (PM-PPr), Cingulo-Opercular (CG-OP), Dorsal Attention-A (dATN-A), Dorsal Attention-B (dATN-B), Salience/Parietal Memory Network (SAL/PMN), Language (LANG), Frontoparietal Network-A (FPN-A), Frontoparietal Network-B (FPN-B), Default Network-A (DN-A) and Default Network-B (DN-B) [32]. These networks can be organized into a three-level architecture: primary networks (VIS-C, VIS-P, SMOT-A, SMOT-B and AUD), intermediate networks (PM-PPr, CG-OP, dATN-A, dATN-B and SAL-PMN) and association networks (LANG, FPN-A, FPN-B, DN-A and DN-B). All cortical entropy aging models at the network level showed decreasing trajectories similar to the global cortical entropy, as shown in Figure 2.

**Fig. 2.**
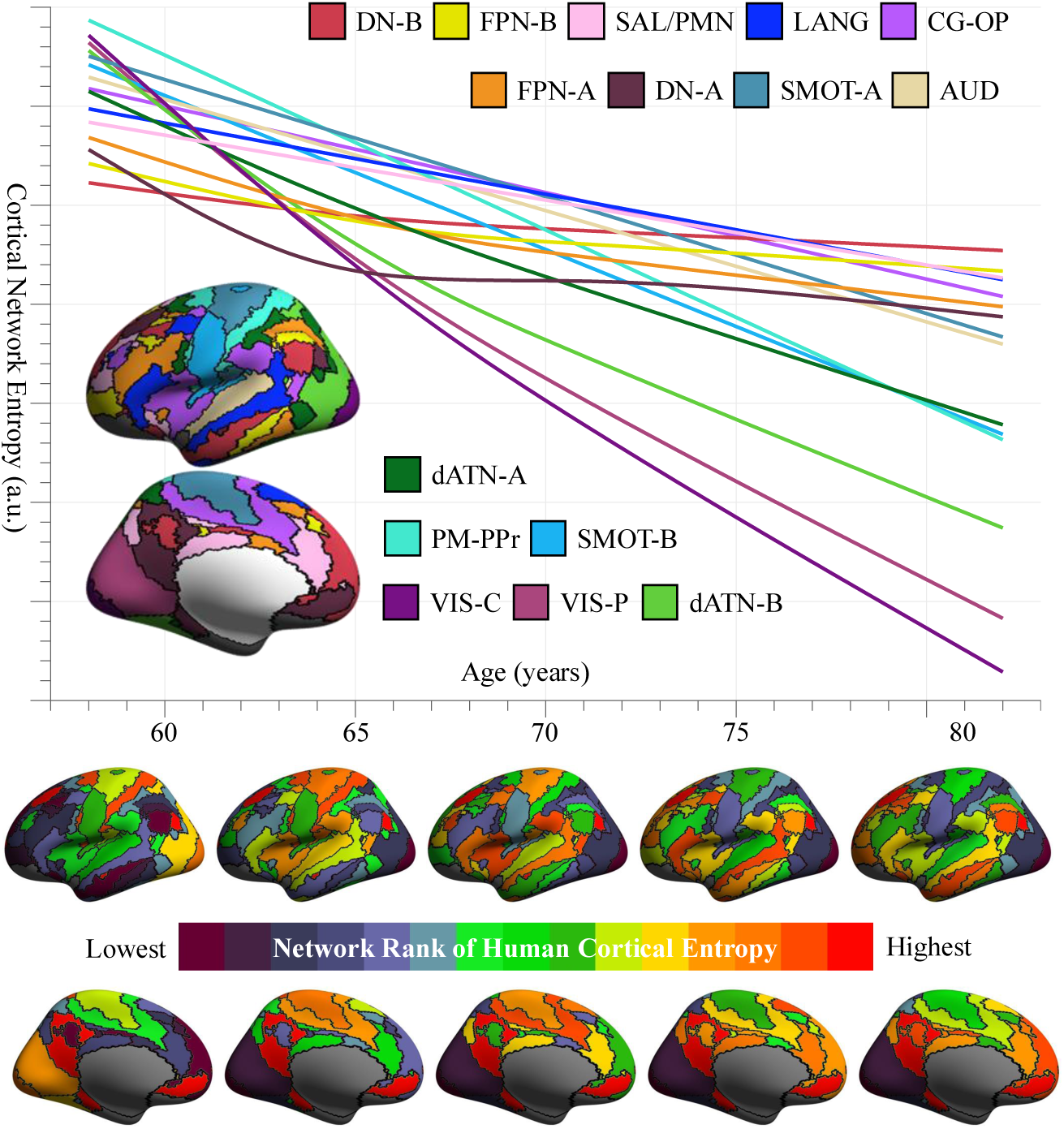
Aging pattern of network entropy in the human cerebral cortex. The fifteen largescale networks of the cerebral cortex are employed to extract their mean entropy values (i.e., cortical network entropy) for trajectory modeling. Samples of each network entropy were separately fitted by GAMM, which estimated the aging curves for individual networks as indicated in the corresponding colors in the DU15NET cortical network parcellation. Accordingly, the rank of complexity level of the fifteen networks from five different ages (58, 64, 70, 76, and 82) were rendered as lateral and medial surfaces at the bottom.

All primary and intermediate networks and LANG (association network) showed a significant (*p <* 0.05) age effect (Table 1). The VIS-C, SMOT-B, PM-PPr, CG-OP, dATN-A and LANG networks showed a significant cohort effect (*p <* 0.05) while the other primary networks (VIS-P and AUD) and intermediate networks (dATN-B and SAL-PMN) only approached the threshold of significance. Higher-order association networks did not show a significant age effect or cohort effect except LANG, but their patterns are similar to the others. For the three-level hierarchy, this aging pattern is clearer: Primary and intermediate areas showed significant effects on age and cohort (*p <* 0.05). The speed of decreasing entropy in primary and intermediate networks is faster than in association networks. That leads to a more interesting observation that the spatial distribution of entropy (the rank of complexity) changes during aging. Specifically, in the early old age, the primary and intermediate networks have a higher entropy than the association networks, but during aging, the default and frontoparietal networks are not significantly affected by aging and the speed of decreasing entropy is much slower, so that in the late old age the association networks have a higher entropy (Figure 2). The diversity among networks also increases with the deepening of aging (Figure 2). This pattern is even clearer within the three-level hierarchy as Figure 3 shows.

**Fig. 3.**
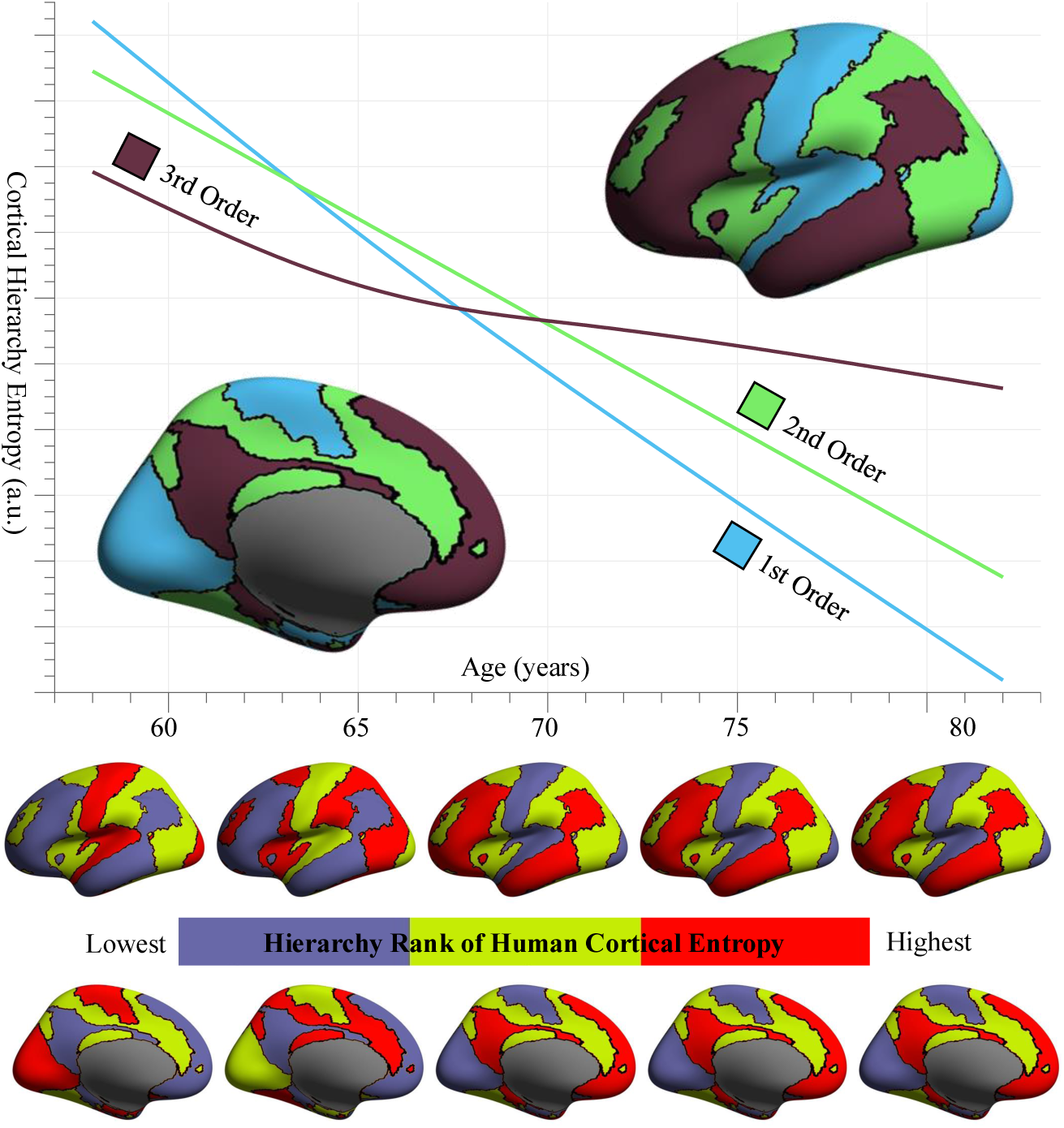
Aging patterns of hierarchical entropy in the human cerebral cortex. A three-level hierarchy of the cerebral cortex is used to extract mean entropy values for fitting the trajectories of primary (the 1st order), intermediate (the 2nd order), and association (the 3rd order) areas by GAMM, which generated the aging curves for the three hierarchies as indicated in the corresponding colors in the DU15NET cortical hierarchical parcellation. Accordingly, the rank of complexity level of the three hierarchies from five different ages (58, 64, 70, 76, and 82) were rendered as lateral and medial surfaces at the bottom.

**Table 1.** Statistical effects on age and cohort of cortical entropy during normal aging. The adjusted *R*^2^ of the GAMM model, *F* –value of age effect, *t*-value of cohort effect and the corresponding *p*-value are listed in the table (∗ *<* 0.05, ∗∗ *<* 0.01, ∗ ∗ ∗ *<* 0.001).

|  | Age (F) | Age (p) | Cohort (t) | Cohort (p) | R-sq. (adj) |
| --- | --- | --- | --- | --- | --- |
| Global | 8.521 | 0.0073 ** | -2.251 | 0.0263 * | 0.169 |
| Primary | 9.830 | 0.0033 ** | -2.110 | 0.0371 * | 0.137 |
| Intermediate | 11.253 | 0.0011 ** | -2.456 | 0.0156 * | 0.143 |
| Association | 1.833 | 0.1390 | -1.519 | 0.1320 | 0.171 |
| VIS-C | 8.956 | 0.0070 ** | -2.039 | 0.0438 * | 0.113 |
| VIS-P | 6.970 | 0.0165 * | -1.913 | 0.0583 | 0.092 |
| SMOT-A | 7.465 | 0.0073 ** | -1.610 | 0.1102 | 0.191 |
| SMOT-B | 9.277 | 0.0029 ** | -2.221 | 0.0283 * | 0.085 |
| AUD | 6.203 | 0.0142 * | -1.935 | 0.0554 | 0.111 |
| PM-PPr | 14.521 | 0.0002 *** | -2.835 | 0.0054 ** | 0.144 |
| CG-OP | 6.326 | 0.0133 * | -2.044 | 0.0432 * | 0.168 |
| dATN-A | 8.898 | 0.0058 ** | -2.430 | 0.0167 * | 0.152 |
| dATN-B | 6.376 | 0.0150 * | -1.875 | 0.0634 | 0.093 |
| SAL-PMN | 4.676 | 0.0327 * | -1.889 | 0.0614 | 0.142 |
| LANG | 4.221 | 0.0425 * | -2.055 | 0.0422 * | 0.169 |
| FPN-A | 3.584 | 0.0713 | -1.595 | 0.1140 | 0.123 |
| FPN-B | 2.070 | 0.2050 | -1.062 | 0.2906 | 0.170 |
| DN-A | 1.613 | 0.1150 | -1.583 | 0.1160 | 0.175 |
| DN-B | 0.900 | 0.5240 | -1.078 | 0.2830 | 0.178 |

### 2.4 Cognitive changes with entropy reduction

We evaluated the longitudinal effect on the performance of the Repeatable Battery for Assessment of Neuropsychological Status (RBANS) [33] to determine whether there are cognitive changes during aging, as described in LongEst2 in 4.6. As shown in Table 2, the total score (*p* = 0.035) and the index scores of immediate memory (*p* = 0.010) and language (*p* = 0.014) exhibited a significant effect on education. This implies that a longer education year is linked to a better test score. No significant effects on age, cohort, genotype, or sex were detectable.

**Table 2.**
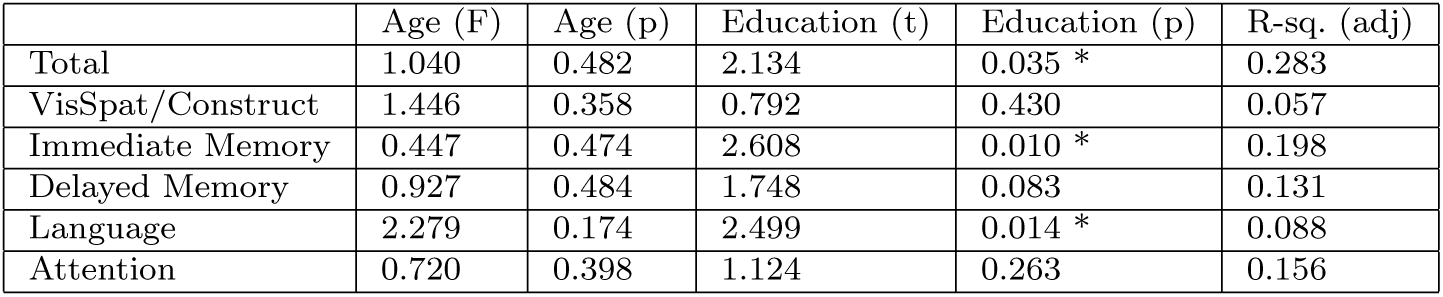
Age and education effects on cognitive performance during aging. The cognitive performance were assessed with the total score and five index scores of RBANS. The adjusted *R*^2^ of the GAMM model, *F* –value of age effect, *t*-value of education effect and the corresponding *p*-value are listed (∗ *<* 0.05).

|  | Age (F) | Age (p) | Education (t) | Education (p) | R-sq. (adj) |
| --- | --- | --- | --- | --- | --- |
| Total | 1.040 | 0.482 | 2.134 | 0.035 * | 0.283 |
| VisSpat/Construct | 1.446 | 0.358 | 0.792 | 0.430 | 0.057 |
| Immediate Memory | 0.447 | 0.474 | 2.608 | 0.010 * | 0.198 |
| Delayed Memory | 0.927 | 0.484 | 1.748 | 0.083 | 0.131 |
| Language | 2.279 | 0.174 | 2.499 | 0.014 * | 0.088 |
| Attention | 0.720 | 0.398 | 1.124 | 0.263 | 0.156 |

The longitudinal change pattern of the RBANS scores with aging cortical entropy was also estimated as described in LongEst3 in 4.6. For the total score, we detected significant entropy effects in the VIS-C (*p* = 0.0463*, R*^2^ = 0.097), SMOT-A (*p* = 0.0262*, R*^2^ = 0.112), SMOT-B (*p* = 0.0274*, R*^2^ = 0.096), PM-PPr (*p* = 0.0240*, R*^2^ = 0.107) and dATN-B (*p* = 0.0440*, R*^2^ = 0.094) networks. This indicated that lower entropy values in these networks lead to higher performance (see Table S1). For the delay memory index, we found significant entropy effects in the SMOT-A (*p* = 0.0065*, R*^2^ = 0.151), SMOT-B (*p* = 0.0051*, R*^2^ = 0.149), PM-PPr (*p* = 0.0099*, R*^2^ = 0.142) and CG-OP (*p* = 0.0342*, R*^2^ = 0.126) and DN-B (*p* = 0.0123*, R*^2^ = 0.144) networks. We also observed significant age effects at baseline. This means that the younger the test participants, the better performance they had (see Table S4). The threelevel hierarchy of the human cortex converged into similar results: significant entropy effects in the primary cortex (*p* = 0.0259*, R*^2^ = 0.107, Table S7) for the total score and significant entropy effects in the primary cortex (*p* = 0.0161*, R*^2^ = 0.133), the intermediate cortex (*p* = 0.0392*, R*^2^ = 0.122) and the global cortex (*p* = 0.0226*, R*^2^ = 0.132, see Table S10). We also observed some significant entropy effects on other index scores; but the adjusted *R*^2^s of these models were too low, indicating a poor model fit.

## 3 Discussion

The present work established cortical entropy trajectories in the human brain during aging and distinguished the effects of age and cohort. Our findings confirmed that cerebral cortex entropy continuously decreases in the aging process both globally and regionally, according to longitudinal data. The entropy of the whole cerebral cortex and most large-scale networks showed such aging effects. These results are consistent with most fMRI studies based on cross-sectional data, as well as studies on other physiological signals such as motor control [34], cardiovascular system [35] and respiratory rhythms[36]. The decreasing of entropy indicates more regular (or, in other words, more predictable) brain activity. For a living system, it is necessary to maintain order (i.e. maintain a relatively low entropy), and temporal coherence is important for the human brain [37, 38]. We are more likely to perceive and act based on experience during aging. The principle of free energy [39] considers the brain as an empirical Bayesian device [40], a living system, to resist disorder and minimize uncertainty (i.e. minimize the free energy). Previous researchers proposed the entropic brain hypothesis [41] that there is a critical zone where normal brain activity occurs, and that normal waking consciousness seems to be placed closer to the order side [42]. Psychedelic compounds shift the brain entropy upward, while sedatives and anesthetics shift the entropy downward. Resting brain entropy of the cerebral cortex has also been found to be negatively correlated with education and task performance in fMRI studies. These results were observed not only in the elderly group [5] but also in adults [4] and adolescents [43]. In the task fMRI study [6], periodic behavior can be reflected in regular brain activity. It has been common to believe that abnormal high or low entropy implies abnormal brain. Our results suggest, at least within a certain range (in the critical zone), for healthy brain, the decreasing entropy means that our brain activity is becoming more order and predictable as interacting with environments. In this respect, older is order.

The cohort effect was found in our study, indicating that individuals born earlier had a higher global entropy. Specifically, the cohort effect is mainly reflected in primary (VIS-C and SMOT-B) and intermediate networks (PM-PPr, CG-OP and dATN-A), with only LANG being detectable among association networks. This effect may even conceal the individual age effect. From a population perspective (cross-sectional), we observed a slightly U-shaped pattern (Figure 1), which may explain why previous studies based on cross-sectional data have reported inconsistent results [5–7, 10–15]. The cohort effect is defined as the effect that having been born in a certain time, region, period or having experienced the same life experience (in the same time period) has on the development or perceptions of a particular group [44]. The environment is one of the main factors that contribute to the effects of the cohort. As mentioned above, fMRI studies have also found a negative correlation between brain entropy and education. These results indicate that, to some extent, the decrease in entropy reflects that brain activity becomes more orderly as individuals interact with the environment and internalize experiences. That might be because the people who were born later lived in a more stable and order society. Our results also suggest that the effect of the environment on the brain might be greater than we previously expected, and future studies (at least for the complexity of low-frequency spontaneous brain activity, as our study revealed) should take the environmental factors into account.

We revealed two specific features in the aging process: the hierarchical shift of the sub-networks and the inter-network diversity of complexity increases during aging. In early old age, cortical complexity ranges from primary unimodal networks to high-order association networks. Networks responsible for primary functions have higher entropy, while networks for high-order functions have relatively lower entropy. A similar pattern has been reported in previous studies, which found that in adults DMN exhibits the lowest entropy in the cerebral cortex [4, 6]. Other researchers have reported quasi-periodic patterns in the default and control networks[45]. The entropy diversity among networks is also low in early old age. Interestingly, this hierarchical organization trend shifts during aging, as the complexity of primary networks decreases more rapidly than that of high-order networks (Figure 3). This implies that primary unimodal networks are more susceptible to the effects of aging, followed by intermediate networks, whereas association networks are not significantly affected by aging, except for the language network. Consequently, in advanced old age, association networks exhibit higher entropy than intermediate and primary networks. The diversity of entropy between networks also increases with aging. In other words, aging is reflected not only in decreasing entropy but also in a shift of the complexity hierarchy of networks, as well as in the diversity of complexity among networks increasing due to different decreasing rates of each network.

Although all subjects had a history of AD from their parents or multiple siblings, the frequency of APOE*ϵ*4 in the sample is not different from the frequency of typical Caucasian populations. Cognitive performance estimation (LongEst2) confirmed that there were no cognitive declines in participants during the 4-year follow-up period. Previous studies have reported that resting brain entropy is negatively correlated with education and general cognitive capacity. In this study, we did not find significant educational effects, which could be due to the fact that the subjects in our study are highly educated (15.08±4.42 years). We also estimated the effect of cognitive performance and found that a lower entropy in VIS-C, SMOT-A, SMOT-B, PM-PPr and dATN-B was associated with a better performance on the total score of RBANS, while a lower entropy in SMOT-A, SMOT-B, PM-PPr, CG-OP and DN-B was associated with a better performance on the delay memory index of RBANS. This might reflect that the RBANS tests contain lots of operations that engage primary and intermediate networks, while as for delay memory high-order association networks are also needed. These findings are consistent with the previously reported negative correlation between entropy and cognitive performance [4].

Our study confirmed that cortical entropy decreases continually in the aging process, both globally and regionally, and revealed its specific features in the aging process: the hierarchical shift of the sub-networks and the internetwork diversity of complexity increases during aging. We conclude the reason why studies based on cross-sectional data have reported inconsistent results: the significant cohort effect. Several limitations must be taken into account. First, although the PREVENT-AD dataset is the best one that is available to us at present for the current study, the sample size is relatively small (24 participants with 120 visits), and in particular there are few very old-age participants. Second, existing data cannot clarify the neurophysiological mechanism underlying the decrease in entropy, and we need more direct evidence in future studies. Third, the typical length of an rsfMRI time series is generally 5 to 10 minutes, which helps reduce the movement and discomfort of the participant. A common time for scanning a single head volume is around 2 seconds, balancing temporal resolution and signal-to-noise ratio. Although a higher temporal resolution would be ideal, previous studies have shown that the entropy metric we used is stable to measure entropy in fMRI time series and the length of the data has only a minor effect on it [4, 6].

## 4 Method

### 4.1 Participants

All participant data included in this study were downloaded from the PREVENT-AD database. It is an open science resource composed of older cognitively unimpaired individuals with a parental or multiple siblings history of AD. The majority of the participants are Caucasian from the greater Montreal area in Qúebec, Canada. There are 24 participants in the observational group (8 males and 16 females, baseline age range: 58-77). They had completed the baseline visit and 4 years of follow-up annually (5 visits each apart from 1 year). During each visit, a standardized cognitive evaluation and MRI scanning session were performed. All participants were cognitively unimpaired during the 4 years follow-up, except one participant who was suspected of probable mild cognitive impairment (MCI) at the last annual visit and this participant is also the one who did not complete all the five RBANS measurements. The individual changing pattern of cortical entropy for this participant is shown in Figure S1.

### 4.2 Imaging parameters

All participants were scanned using a Siemens TIM Trio 3 Tesla MRI scanner with a Siemens standard 12 channel coil (Siemens Medical Solutions, Erlangen, Germany) at the Brain Imaging Centre of the Douglas Mental Health University Institute. During each visit, a structural MRI and two rsfMRI scans were acquired. Structural MRI images were acquired using a T1-weighted magnetization prepared gradient echo (MPRAGE) sequence (3D sagittal view, TR= 2300*ms*, TE= 2.98*ms*, TI= 900*ms*, flip angle 9^◦^, FOV= 256 240 176*mm*, phase encode A-P, BW= 240*Hz/px*, GRAPPA 2, Resolution= 1 1 1*mm*^3^, Scan time= 5.12*min*). RsfMRI data were acquired with an echo-planar imaging (EPI) sequence (with following parameters: 2D axial view, TR= 2000*ms*; TE= 30*ms*; flip angle= 90^◦^, FOV= 256 *×* 256*mm*, 32 slices, phase encode A-P, BW= 2442*/px*, resolution= 4 *×* 4 *×* 4*mm*^3^, scan time= 5.04*min*).

### 4.3 Cognitive evaluation

Cognitive performance was assessed using the RBANS at baseline and each subsequent follow-up visit. The battery consists of 12 subtests and yields 5 index scores (immediate memory, delayed memory, language, attention and visuospatial capacities) as well as a total score. It is available in both French and English in 4 equivalent versions to reduce practice effects in longitudinal assessment. For these participants, the versions were administered in chronologic order A, B, C, D. The data were scored following the RBANS manual, with two versions of scores provided. One is the age-adjusted index scores, the other scored all participants using norms specified for individuals aged 60-69 years which allows detection of potential changes with advancing age. In our study, we used the 60-69 age norm scores.

### 4.4 Image data preprocessing

Anatomical T1 images were visually inspected and then uploaded to the volBrain pipeline (http://volbrain.upv.es) [46] for noise removal, bias correction, intensity normalization, and brain extraction. All brain extractions underwent visual inspection to ensure tissue integrity. After initial quality checks, the T1 images were passed into the Connectome Computation System (https://github.com/zuoxinian/CCS) [47] for surface-based analyses.

RsfMRI data were preprocessed following steps: (1) dropping the first 10s (five TRs) for the equilibrium of the magnetic field; (2) head motion correction; (3) slicing timing; (4) de-spiking the time series; (5) estimating head motion parameters; (6) registering functional images to high resolution T1 images using boundary-based registration; (7) mitigating nuisance effects such as ICA-AROMA-derived [48], CSF and white matter signals; (8) removing linear and quadratic trends of the time series; (9) projecting volumetric time series to both native surface space and fsaverage5 surface space. All preprocessing scripts of the above steps are available on github (https://github.com/zuoxinian/CCS/tree/master/H1) [49].

### 4.5 Entropy calculation

Cortical entropy has been reported to be significantly lower than that of white matter and subcortical structures [4, 6]. Thus, we focus solely on the cerebral cortex. The entropy value was calculated at each vertex based on the preprocessed rsfMRI time series with BENtbx (https://cfn.upenn.edu/~zewang/ BENtbx.php) by using an approximate entropy measurement (SampEn) with the parameters recommended in [6]. This measure is an improved metric of approximate entropy [50]. SampEn is based on the entropy of measured hemodynamic states, which considers dependency over time using temporal embedding. In other words, it reflects the statistical dependencies or order implicit in itinerant dynamics expressed over extended periods of time. Specifically, for the preprocessed BOLD time series at one vertex, denoted by *x* = [*x*_1_*, x*_2_*, …, x*_N_], where *N* is the number of time points. With a predefined dimension *m* and a distance threshold *r* (we use *m* = 3 and *r* is the 0.6 times the standard deviation of *x*), we extracted a series of embedded vectors from *x*, each with *m* consecutive points: *u_i_* = [*x_i_, x_i_* + 1*, …, x_i_* + *m* 1], where 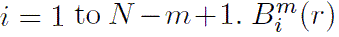 counts the number of *u_j_* (*j* = 1 to *N − m*, and *j* ≠ *i*) whose Chebyshev distance to *u_i_* is less than *r*, as does 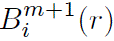 for the dimension of *m* + 1. By averaging across all possible vectors, we have (as shown in [6]):

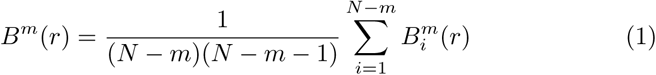

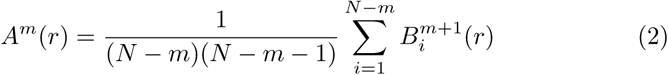

SampEn is calculated as:

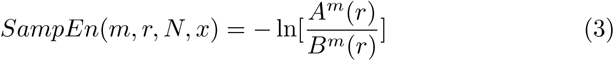

The global entropy was calculated based on the data in native surface space. For each processed rsfMRI data, the mean entropy of the whole cerebral cortex was calculated. Since each participant underwent two rsfMRI scans at each visit, we averaged the two mean entropy values to obtain the global entropy for each visit, used for further analysis. We also calculated the mean entropy values of fifteen networks and three hierarchies as documented in [32]. Specifically, the three-level hierarchy is primary, intermediate, and association. Each of them consist of five networks: primary networks include VIS-C, VIS-P, SMOT-A, SMOT-B and AUD; intermediate networks include PM-PPr, CGOP, dATN-A, dATN-B and SAL-PMN; association networks include LANG, FPN-A, FPN-B, DN-A and DN-B.

### 4.6 Statistical modeling

We use GAMM to estimate the global entropy trajectory and the mean entropy values of 15 networks and three hierarchies separately during aging and distinguish the effects of age and cohort. We modeled the effects of age on entropy with smooth functions constructed as weighted sums of *k* basis functions. In this study, we employed cubic regression splines and chose *k* = 5 as large enough to allow a wide range of nonlinear patterns to be estimated, while small enough to minimize overfitting and allow computational efficiency. Random effect was included for longitudinal data with repeated measurements. The linear age-independent cohort effect was modeled by including the birth date. Since education [4, 5] and sex effects of resting brain entropy have been reported in previous studies [4], we also took these factors into account in GAMM. Additionally, the gene effect was taken into account in our model, not only the main effect but also its interaction with age estimated by smooth functions. This is due to the fact that the APOE*ϵ*4 allele is a well-known risk genotype for AD and our data have a parental or multiple-sibling history of AD. The specific model was constructed on the basis of the recommendation in [51] using the gamm4 package (https://CRAN.R-project.org/package=gamm4).

Although the integrity of cognition of our participants was confirmed by the Montreal Cognitive Assessment [52] and the Clinical Dementia Rating Scale [53] at the eligibility visit and by RBANS at each visit, the effect of age on the performance of RBANS was evaluated to find out whether there are cognitive declines during aging. The total score and the five index scores were estimated using a model similar to that mentioned previously. One participant who didn’t complete all the five cognitive tests was excluded from the analysis. This participant was also suspected of probable MCI at his last annual visit as described in 4.1. Furthermore, the pattern of longitudinal changes of the RBANS scores with entropy during aging was explored. Because both variables have the same effects of sex, genotype and education years, so in this model, despite using the smooth functions to model the effect of entropy, we only considered the fixed effect of the age at initial visit and the random effect.

## Declarations

### Ethics approval and consent to participate

Protocols, consent forms and study procedures were approved by McGill Institutional Review Board and/or Douglas Mental Health University Institute Research Ethics Board. Specific consent forms were presented prior to each experimental procedure. More details can be found in [29].

### Consent for publication

Participants have obtained informed consent to publicly share their anonymized data or publish the data in an open online publication.

### Availability of data and materials

All data included in this study can be downloaded from the PREVENT-AD database (https://registeredpreventad.loris.ca). All codes in this study can be requested from the authors.

### Authors’ contributions

DC worked on data curation, design of models, software, validation, visualization, and preparation of the original draft. XNZ and XW have contributed to conceptualization, formal analysis, design of models, supervision, validation, and editing of the draft. All authors have revised the paper.

## Supplementary information

**Figure S1.**
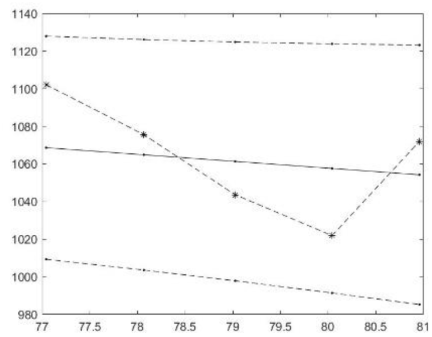
The individual BEN changing pattern during the four years of the specific subject: the dot-dash line represents the real global BEN changing pattern, and the solid line represents the fitted trajectory, the two dash lines are the 95% confidence interval.

Longitudinal changing pattern of the RBANS scores with BEN during aging

**Fifteen networks**

**Table S1.** BEN and baseline-age effects of cognitive performance during aging, total score of RBANS.

|  | BEN (F) | BEN (p) | Age-bl (t) | Age-bl (p) | R-sq. (adj) |
| --- | --- | --- | --- | --- | --- |
| VIS-C | 4.061 | 0.0463 * | -1.684 | 0.0951 | 0.097 |
| VIS-P | 3.635 | 0.0592 | -1.693 | 0.0932 | 0.095 |
| SMOT-A | 5.075 | 0.0262 * | -1.814 | 0.0723 | 0.112 |
| SMOT-B | 4.995 | 0.0274 * | -1.675 | 0.0967 | 0.096 |
| AUD | 2.272 | 0.2160 | -1.632 | 0.1060 | 0.068 |
| PM-PPr | 5.238 | 0.0240 * | -1.715 | 0.0890 | 0.107 |
| CG-OP | 1.933 | 0.1790 | -1.610 | 0.1100 | 0.091 |
| dATN-A | 2.083 | 0.1520 | -1.674 | 0.0968 | 0.095 |
| dATN-B | 4.149 | 0.0440 * | -1.678 | 0.0962 | 0.094 |
| SAL/PMN | 0.641 | 0.5530 | -1.623 | 0.1070 | 0.077 |
| LANG | 2.028 | 0.2790 | -1.576 | 0.1180 | 0.087 |
| FPN-A | 3.640 | 0.0725 | -1.635 | 0.1050 | 0.094 |
| FPN-B | 1.721 | 0.2340 | -1.641 | 0.1040 | 0.091 |
| DN-A | 0.494 | 0.4840 | -1.569 | 0.1200 | 0.081 |
| DN-B | 0.602 | 0.4400 | -1.579 | 0.1170 | 0.088 |

This table shows the BEN effect of the total score of RBANS for each brain network, the effect of the age that participants took part in the initial MRI session (baseline) were also estimated. The adjusted $R^2$ of the model, F-value of BEN effect, t-value of baseline-age effect and the corresponding $p$ -value are listed in the table.

**Table S2.** BEN and baseline-age effects of cognitive performance during aging, visuospatial/constructional score.

|  | BEN (F) | BEN (p) | Age-bl (t) | Age-bl (p) | R-sq. (adj) |
| --- | --- | --- | --- | --- | --- |
| VIS-C | 5.359 | 0.0225 * | -0.915 | 0.3622 | 0.0129 |
| VIS-P | 4.514 | 0.0358 * | -0.900 | 0.3702 | -0.006 |
| SMOT-A | 3.993 | 0.0481 * | -1.000 | 0.3196 | -0.001 |
| SMOT-B | 4.428 | 0.0263 * | -0.869 | 0.3866 | 0.008 |
| AUD | 1.976 | 0.1630 | -0.819 | 0.4144 | 0.002 |
| PM-PPr | 4.315 | 0.0430 * | -0.912 | 0.3637 | 0.011 |
| CG-OP | 4.250 | 0.0416 * | -0.754 | 0.4526 | 0.008 |
| dATN-A | 6.066 | 0.0153 * | -0.898 | 0.3713 | 0.014 |
| dATN-B | 3.031 | 0.1190 | -0.905 | 0.3675 | 0.010 |
| SAL/PMN | 3.634 | 0.0874 | -0.747 | 0.4569 | 0.023 |
| LANG | 3.521 | 0.0632 | -0.742 | 0.4598 | 0.022 |
| FPN-A | 2.967 | 0.0533 | -0.814 | 0.4171 | 0.030 |
| FPN-B | 3.212 | 0.1110 | -0.788 | 0.4323 | 0.042 |
| DN-A | 4.067 | 0.0461 * | -0.635 | 0.5267 | 0.006 |
| DN-B | 3.975 | 0.0432 * | -0.644 | 0.5208 | 0.030 |

This table shows the BEN effect of the visuospatial/constructional score of RBANS for each brain network, the effect of the age that participants took part in the initial MRI session (baseline) were also estimated. The adjusted $R^2$ of the model, F-value of BEN effect, t-value of baseline-age effect and the corresponding $p$ -value are listed in the table.

**Table S3.** BEN and baseline-age effects of cognitive performance during aging, immediate memory score.

|  | BEN (F) | BEN (p) | Age-bl (t) | Age-bl (p) | R-sq. (adj) |
| --- | --- | --- | --- | --- | --- |
| VIS-C | 0.086 | 0.769 | -0.727 | 0.469 | 0.00229 |
| VIS-P | 0.138 | 0.711 | -0.816 | 0.416 | 0.00785 |
| SMOT-A | 0.978 | 0.501 | -0.907 | 0.367 | 0.0347 |
| SMOT-B | 0.396 | 0.53 | -0.823 | 0.412 | 0.0162 |
| AUD | 0.175 | 0.76 | -0.807 | 0.421 | -0.0147 |
| PM-PPr | 0.914 | 0.341 | -0.858 | 0.393 | 0.0268 |
| CG-OP | 0.861 | 0.418 | -0.786 | 0.434 | -0.000807 |
| dATN-A | 0.014 | 0.904 | -0.798 | 0.427 | 0.00165 |
| dATN-B | 0.738 | 0.392 | -0.835 | 0.405 | 0.0189 |
| SAL/PMN | 0.7 | 0.405 | -0.814 | 0.418 | -0.0343 |
| LANG | 3.167 | 0.0921 | -0.751 | 0.454 | -0.00727 |
| FPN-A | 1.871 | 0.215 | -0.742 | 0.46 | 0.0354 |
| FPN-B | 2.44 | 0.0452 * | -0.643 | 0.521 | 0.0323 |
| DN-A | 1.743 | 0.189 | -0.908 | 0.366 | -0.0404 |
| DN-B | 0.728 | 0.395 | -0.853 | 0.396 | -0.0297 |

This table shows the BEN effect of the immediate memory score of RBANS for each brain network, the effect of the age that participants took part in the initial MRI session (baseline) were also estimated. The adjusted $R^2$ of the model, F-value of BEN effect, t-value of baseline-age effect and the corresponding $p$ -value are listed in the table.

**Table S4.**
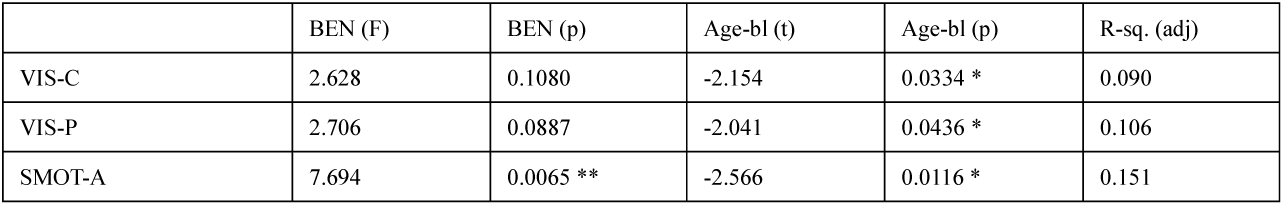
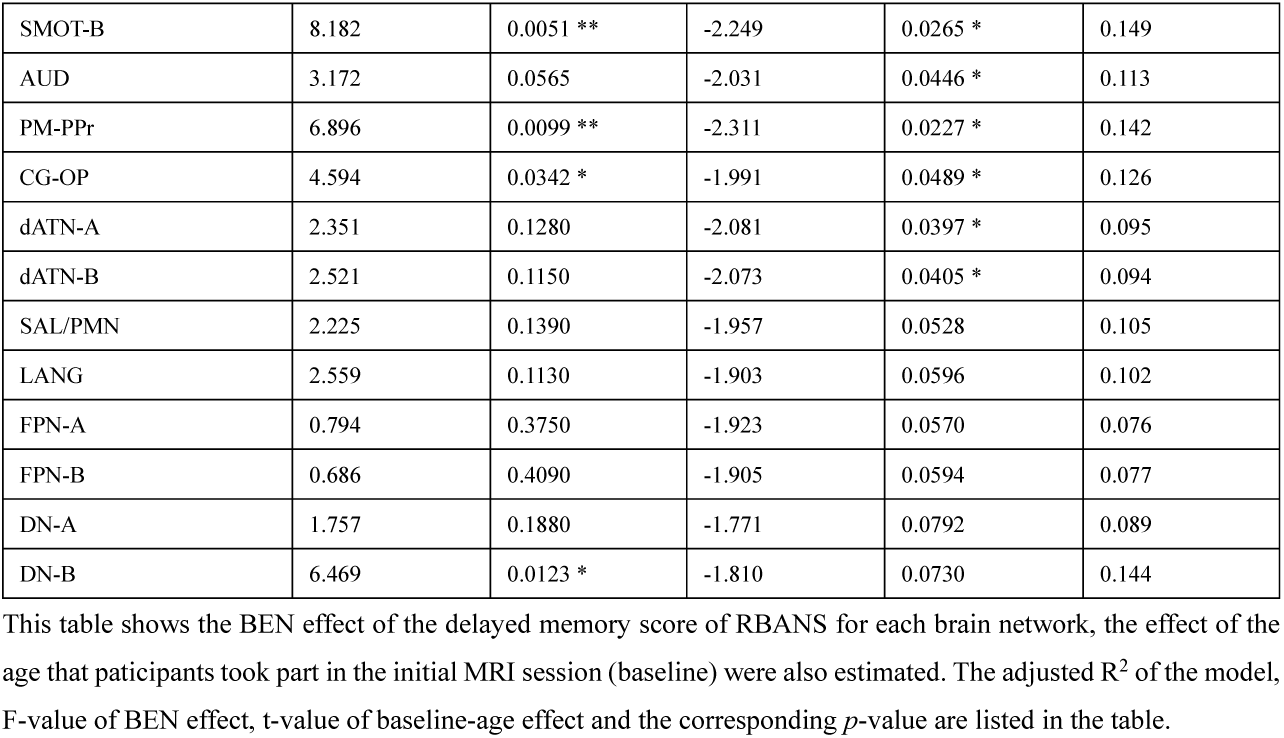
BEN and baseline-age effects of cognitive performance during aging, delayed memory score.

**Table S5.** BEN and baseline-age effects of cognitive performance during aging, language score.

|  | BEN (F) | BEN (p) | Age-bl (t) | Age-bl (p) | R-sq. (adj) |
| --- | --- | --- | --- | --- | --- |
| VIS-C | 0.554 | 0.4580 | -0.929 | 0.3550 | -0.001 |
| VIS-P | 0.205 | 0.6510 | -0.906 | 0.3670 | -0.001 |
| SMOT-A | 0.501 | 0.4810 | -0.978 | 0.3300 | 0.006 |
| SMOT-B | 0.216 | 0.6430 | -0.901 | 0.3690 | 0.002 |
| AUD | 0.170 | 0.6810 | -0.878 | 0.3820 | 0.003 |
| PM-PPtr | 0.636 | 0.4270 | -0.938 | 0.3500 | 0.008 |
| CG-OP | 0.347 | 0.7510 | -0.877 | 0.3820 | 0.009 |
| dATN-A | 0.267 | 0.6070 | -0.904 | 0.3680 | 0.002 |
| dATN-B | 0.639 | 0.4260 | -0.912 | 0.3640 | 0.002 |
| 5SAL/PMN | 0.045 | 0.9500 | -0.868 | 0.3870 | -0.001 |
| LANG | 1.614 | 0.3600 | -0.781 | 0.4360 | 0.017 |
| FPN-A | 3.001 | 0.0243 * | -0.848 | 0.3980 | 0.067 |
| FPN-B | 0.036 | 0.8490 | -0.860 | -0.3920 | -0.002 |
| DN-A | 0.015 | 0.9040 | -0.846 | 0.3990 | -0.004 |
| DN-B | 0.027 | 0.8700 | -0.848 | 0.3980 | -0.002 |

This table shows the BEN effect of the language score of RBANS for each brain network, the effect of the age that participants took part in the initial MRI session (baseline) were also estimated. The adjusted $R^2$ of the model, F-value of BEN effect, t-value of baseline-age effect and the corresponding $p$ -value are listed in the table.

**Table S6.** BEN and baseline-age effects of cognitive performance during aging, attention score.

|  | BEN (F) | BEN (p) | Age-bl (t) | Age-bl (p) | R-sq. (adj) |
| --- | --- | --- | --- | --- | --- |
| VIS-C | 1.743 | 0.1890 | -1.513 | 0.1332 | 0.067 |
| VIS-P | 2.548 | 0.1130 | -1.580 | 0.1169 | 0.066 |
| SMOT-A | 4.501 | 0.0361 * | -1.659 | 0.1000 | 0.057 |
| SMOT-B | 2.979 | 0.0871 | -1.550 | 0.1239 | 0.053 |
| AUD | 3.131 | 0.0795 | -1.512 | 0.1333 | 0.061 |
| PM-PPr | 2.729 | 0.101 | -1.574 | 0.1184 | 0.059 |
| CG-OP | 1.953 | 0.1650 | -1.492 | 0.1386 | 0.063 |
| dATN-A | 1.242 | 0.2680 | -1.559 | 0.1218 | 0.063 |
| dATN-B | 2.001 | 0.1600 | -1.562 | 0.1210 | 0.062 |
| SAL/PMN | 1.074 | 0.3020 | -1.497 | 0.1372 | 0.065 |
| LANG | 2.427 | 0.1220 | -1.468 | 0.1450 | 0.062 |
| FPN-A | 1.298 | 0.3900 | -1.523 | 0.1305 | 0.066 |
| FPN-B | 0.132 | 0.8860 | -1.545 | 0.1253 | 0.068 |
| DN-A | 2.128 | 0.1470 | -1.417 | 0.1593 | 0.057 |
| DN-B | 1.136 | 0.2890 | -1.465 | 0.1458 | 0.064 |

This table shows the BEN effect of the attention score of RBANS for each brain network, the effect of the age that participants took part in the initial MRI session (baseline) were also estimated. The adjusted $R^2$ of the model, F-value of BEN effect, t-value of baseline-age effect and the corresponding $p$ -value are listed in the table.

**Global and three-level hierarchy**

**Table S7.** BEN and baseline-age effects of cognitive performance during aging, total score of RBANS.

|  | BEN (F) | BEN (p) | Age-bl (t) | Age-bl (p) | R-sq. (adj) |
| --- | --- | --- | --- | --- | --- |
| Global | 2.849 | 0.0942 | -1.645 | 0.1030 | 0.101 |
| Primary | 5.098 | 0.0259 * | -1.738 | 0.0849 | 0.107 |
| Intermediate | 2.920 | 0.0903 | -1.650 | 0.1020 | 0.102 |
| Association | 1.284 | 0.3410 | -1.578 | 0.1170 | 0.091 |

This table shows the BEN effect of the attention score of RBANS for global and each hierarchy, the effect of the age that participants took part in the initial MRI session (baseline) were also estimated. The adjusted $R^2$ of the model, F-value of BEN effect, t-value of baseline-age effect and the corresponding $p$ -value are listed in the table.

**Table S8.**
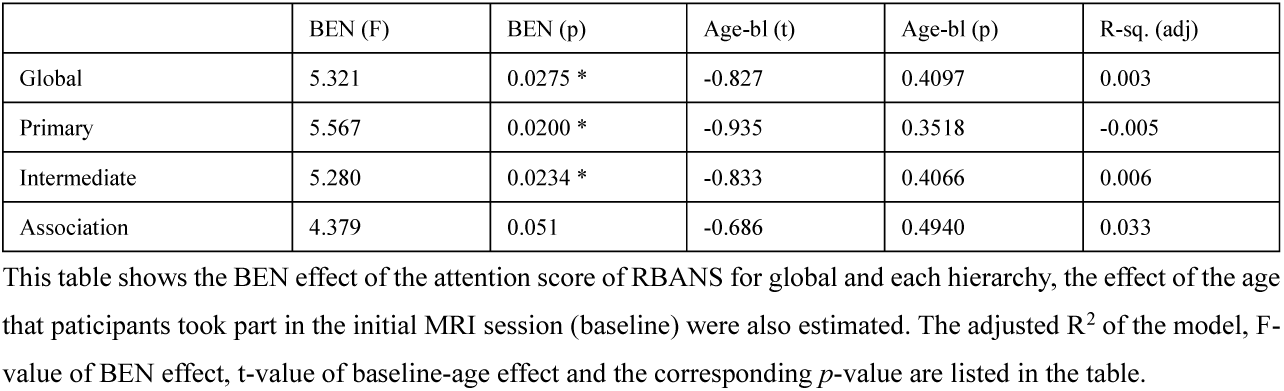
BEN and baseline-age effects of cognitive performance during aging, visuospatial/constructional score.

|  | BEN (F) | BEN (p) | Age-bl (t) | Age-bl (p) | R-sq. (adj) |
| --- | --- | --- | --- | --- | --- |
| Global | 5.321 | 0.0275 * | -0.827 | 0.4097 | 0.003 |
| Primary | 5.567 | 0.0200 * | -0.935 | 0.3518 | -0.005 |
| Intermediate | 5.280 | 0.0234 * | -0.833 | 0.4066 | 0.006 |
| Association | 4.379 | 0.051 | -0.686 | 0.4940 | 0.033 |

This table shows the BEN effect of the attention score of RBANS for global and each hierarchy, the effect of the age that participants took part in the initial MRI session (baseline) were also estimated. The adjusted $R^2$ of the model, F-value of BEN effect, t-value of baseline-age effect and the corresponding $p$ -value are listed in the table.

**Table S9.**
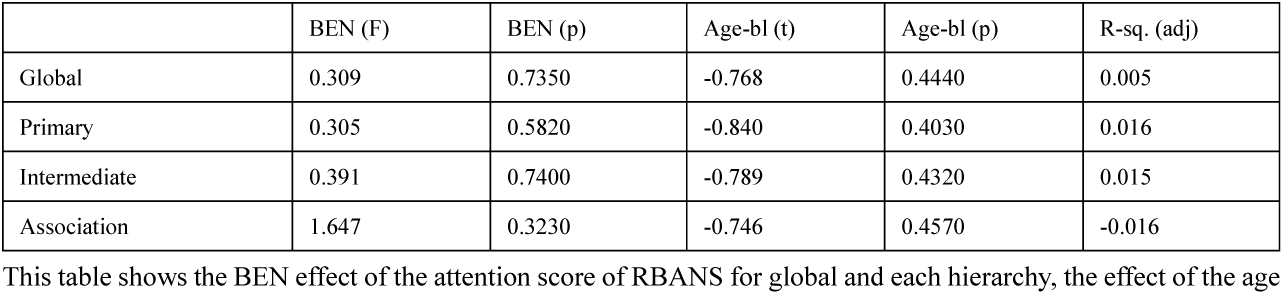
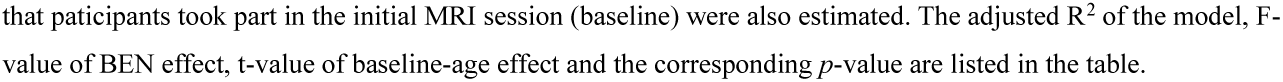
BEN and baseline-age effects of cognitive performance during aging, immediate memory score.

**Table S10.** BEN and baseline-age effects of cognitive performance during aging, delayed memory score.

|  | BEN (F) | BEN (p) | Age-bl (t) | Age-bl (p) | R-sq. (adj) |
| --- | --- | --- | --- | --- | --- |
| Global | 5.350 | 0.0226 * | -2.137 | 0.0348 * | 0.132 |
| Primary | 5.971 | 0.0161 * | -2.313 | 0.0226 * | 0.133 |
| Intermediate | 4.355 | 0.0392 * | -2.116 | 0.0365 * | 0.122 |
| Association | 2.821 | 0.0958 | -1.868 | 0.0643 | 0.108 |

This table shows the BEN effect of the attention score of RBANS for global and each hierarchy, the effect of the age that participants took part in the initial MRI session (baseline) were also estimated. The adjusted $R^2$ of the model, F-value of BEN effect, t-value of baseline-age effect and the corresponding $p$ -value are listed in the table.

**Table S11.** BEN and baseline-age effects of cognitive performance during aging, language score.

|  | BEN (F) | BEN (p) | Age-bl (t) | Age-bl (p) | R-sq. (adj) |
| --- | --- | --- | --- | --- | --- |
| Global | 0.282 | 0.5960 | -0.883 | 0.3790 | 0.003 |
| Primary | 0.386 | 0.5360 | -0.931 | 0.3540 | 0.004 |
| Intermediate | 0.354 | 0.5530 | -0.886 | 0.3770 | 0.004 |
| Association | 0.380 | 0.8220 | -0.787 | 0.4330 | 0.008 |

This table shows the BEN effect of the attention score of RBANS for global and each hierarchy, the effect of the age that participants took part in the initial MRI session (baseline) were also estimated. The adjusted $R^2$ of the model, F-value of BEN effect, t-value of baseline-age effect and the corresponding $p$ -value are listed in the table.

**Table S12.** BEN and baseline-age effects of cognitive performance during aging, attention score.

|  | BEN (F) | BEN (p) | Age-bl (t) | Age-bl (p) | R-sq. (adj) |
| --- | --- | --- | --- | --- | --- |
| Global | 2.830 | 0.0953 | -1.525 | 0.1301 | 0.058 |
| Primary | 4.103 | 0.0452 * | -1.600 | 0.1125 | 0.060 |
| Intermediate | 2.433 | 0.1220 | -1.531 | 0.1285 | 0.060 |
| Association | 1.491 | 0.2250 | -1.467 | 0.1453 | 0.064 |

This table shows the BEN effect of the attention score of RBANS for global and each hierarchy, the effect of the age that participants took part in the initial MRI session (baseline) were also estimated. The adjusted $R^2$ of the model, F-value of BEN effect, t-value of baseline-age effect and the corresponding $p$ -value are listed in the table.

